# Chronic BDNF simultaneously inhibits and unmasks superficial dorsal horn neuronal activity

**DOI:** 10.1101/2020.11.02.364786

**Authors:** Sascha R.A. Alles, Max A. Odem, Van B. Lu, Ryan M. Cassidy, Peter A. Smith

## Abstract

Brain-derived neurotrophic factor (BDNF) is critically involved in the pathophysiology of chronic pain. However, the mechanisms of BDNF action on specific neuronal populations in the spinal superficial dorsal horn (SDH) requires further study. We used chronic BDNF treatment (200 ng/ml, 5-6 days) of defined-medium, serum-free spinal organotypic cultures to study intracellular calcium ([Ca2+]_i_) fluctuations. A detailed quantitative analysis of these fluctuations using the Frequency-independent biological signal identification (FIBSI) program revealed that BDNF simultaneously depressed activity in some SDH neurons while it unmasked a particular subpopulation of ‘silent’ neurons causing them to become spontaneously active. Blockade of gap junctions disinhibited a subpopulation of SDH neurons and reduced BDNF-induced synchrony in BDNF-treated cultures. BDNF reduced neuronal excitability by measuring spontaneous excitatory postsynaptic currents. This was similar to the depressive effect of BDNF on the [Ca2+]_i_ fluctuations. This study reveals novel regulatory mechanisms of SDH neuronal excitability in response to BDNF.

## Introduction

Injury to, or disease of, the somatosensory system frequently generates chronic and sometimes intractable neuropathic pain (1,2). In experimental animals, peripheral nerve damage, such as that generated by chronic constriction or section of the sciatic nerve, induces pain-related behaviours that serve as a model for human neuropathic pain (3,4). Seven or more days of sciatic nerve injury promote an enduring increase in the excitability of first order primary afferent neurons (5–8). These become chronically active and release a variety of mediators (cytokines, chemokines, neuropeptides, ATP and growth factors) that predispose spinal microglia to a more ‘activated’ state (9–14). These in turn, release further mediators, including brain derived neurotrophic factor (BDNF) that promote a slowly developing, but persistent increase in excitability of second order neurons in the spinal dorsal horn. This ‘central sensitization’ is thought to be responsible for the allodynia, hyperalgesia, spontaneous pain and causalgia that characterize neuropathic pain (3,15). Spinal actions of BDNF involve alteration in Cl^-^ gradients such that the normally inhibitory actions of GABA become excitatory (16,17). There is also increased excitatory synaptic drive to putative excitatory neurons (9,18). This results in an overall increase in excitability as monitored by confocal Ca^2+^ imaging in organotypic cultures of rat or mouse spinal cord (9,18–20). Despite this, the long-term effects of upregulated BDNF on neuronal plasticity in the superficial dorsal horn (SDH) are not fully understood.

Naïve cultures display spontaneous oscillatory changes in intracellular Ca^2+^ levels [Ca2+]_i_ and we have previously shown that the amplitude and frequency of these changes in [Ca2+]_i_ are profoundly increased following 5-6 d treatment with BDNF (20). We used this model and a new Frequency-independent biological signal identification (FIBSI) (21) program to quantitatively measure the fluctuations of [Ca2+]_i_ and examine their synchronicity. Unexpectedly, we observed two opposite effects of BDNF that appeared to occur simultaneously. First, BDNF caused an overall decrease in fluctuation size. Second, we noticed a particular population of SDH neurons in naïve cultures that did not display typical, marked fluctuations of [Ca2+]_i_. This population of ‘silent’ neurons was absent in BDNF-treated cultures, suggesting that BDNF unmasks these ‘silent’ neurons and causes them to become spontaneously active. We next investigated the role of gap junctions in mediating the [Ca2+]_i_ fluctuations; application of the gap junction blocker octanol to chronic BDNF-treated neurons revealed a subpopulation of neurons that generate low-frequency, large [Ca2+]_i_ fluctuations. Further pharmacological experiments indicated the [Ca2+]_i_ fluctuations in active neurons are regulated by diverse mechanisms including voltage-gated calcium and sodium channels as well as GABA and NMDA receptors. Finally, we used FIBSI to reanalyze a previous dataset of spontaneous excitatory postsynaptic current (sEPSC) recordings from BDNF-treated dorsal horn neurons in order to qualitatively compare the effects of BDNF on the sEPSCs and [Ca2+]_i_ fluctuations in SDH neurons. The depressive effects of BDNF on the sEPSCs in delay and tonic-firing SDH neurons were consistent with the effects on the [Ca2+]_i_ fluctuations. This study reveals novel mechanisms of BDNF regulation of dorsal horn excitability, which has implications for the study of chronic neuropathic pain physiology.

## Results

### Chronic BDNF treatment of spinal organotypic cultures simultaneously depresses and unmasks [Ca2+]_i_ fluctuations in superficial dorsal horn neurons

In previous work we identified that chronic BDNF treatment (200 ng/ml, 5-6 days) induces fluctuations compared to naïve cultures (20,22). In this study, we performed a detailed quantitative analysis of the fluctuations in [Ca2+]_i_ using Fluo-4 AM Ca2+ imaging and the FIBSI analysis program (21) to further establish the nature of BDNF-induced changes in lamina II neurons of the SDH. Refer to **Fig 1A1-A3** for an example of [Ca2+]_i_ fluctuations detected by the FIBSI program. Inspection of the FIBSI-processed recordings revealed a subset of naïve, untreated neurons were ‘silent’ (**Fig 1B**, left) for the duration of the recording with notably smaller amplitudes (‘silent’ = 1.7 ± 0.04 AU; naïve = 53.7 ± 1.6 AU; BDNF = 25.5 ± 0.7 AU). No ‘silent’ neurons were found in the BDNF treatment group (**Fig 1B**, right). Refer to **Table 1** for detailed statistical information. This stark contrast suggested BDNF may be unmasking, or activating, these ‘silent’ neurons. We grouped the fluctuations in the ‘silent’ neurons together for further comparison to the fluctuations in the other active naïve neurons and BDNF-treated neurons. Log-transformed fluctuation amplitudes (**Fig 1C**) and area under the curve (AUC; **Fig 1D**) were larger in the naïve and BDNF conditions compared to ‘silent’. Unexpectedly, both measures were significantly decreased in the BDNF condition compared to untreated. A plausible explanation for this effect was that chronic BDNF induced a concomitant enhancement of [Ca2+]_i_ fluctuations in some neurons and depression in others. Indeed, the effects of BDNF on the relative frequencies (%) for the fluctuation amplitudes (amplitudes were normalized to the largest fluctuation in each neuron) revealed a biphasic effect; frequencies of the smaller- and larger-amplitude fluctuations were greater in the BDNF-treated neurons compared to the naïve and ‘silent’ neurons (**Fig 1E**). We next compared the mean parameters between neurons in order to control for differences in sampling periods. A <10% maximal fluctuation amplitude cut-off was applied to each neuron to reduce the effects of low-amplitude noise. As expected, nonparametric comparisons indicated that fluctuation amplitude (**Fig 1F**) and AUC (**Fig 1G**) were both greater in the naïve and BDNF-treated neurons compared to the ‘silent’ neurons, but were not significantly different between naïve and BDNF-treated neurons. Further analysis of the fluctuation kinetics indicated no effect on duration (**Fig 1H**), a modest, but significant decrease in rise time (i.e., peak time is more negative) compared to the ‘silent’ neurons (**Fig 1I**), and no effect on frequency (**Fig 1J**).

**Figure 1.**
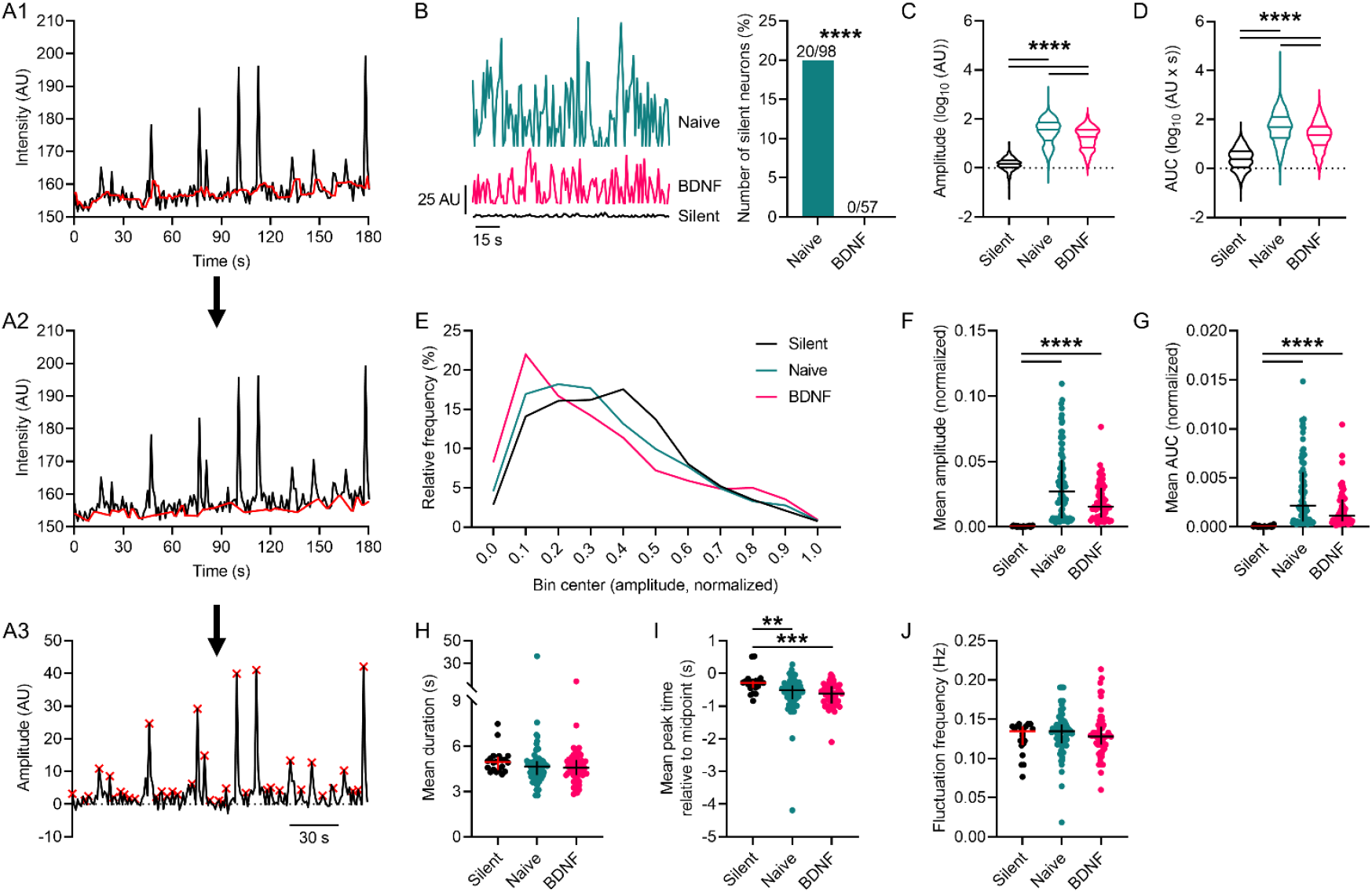
BDNF-induced fluctuations of [Ca2+]_I_ in superficial dorsal horn neurons. **(A1-A3)** Example recording [Ca2+]_i_ fluctuations sampled in a SDH neuron and the processing steps used by the FIBSI event-detection program. The running median (red line in A1, window = ∼5 s) was calculated based on the raw amplitude (AU) and time coordinate data, then the peaks below the running median were traced to form a reference line (red line in A2), and then the Ramer-Douglas-Peucker algorithm detected waveforms above the reference (red x at event peaks). **(B)** Left: Examples of [Ca2+]_i_ activity detected by FIBSI in 3 different neurons. Right: The proportion of ‘silent’ neurons in the BDNF-treatment condition was significantly reduced compared to the naïve condition. Proportions were compared using a Fisher exact test. **(C-D)** Violin plots of the log-transformed values for fluctuation amplitude and AUC. Fluctuation amplitudes and AUC were significantly increased in naïve and BDNF-treated neurons compared to the ‘silent’ neurons, while both measures in the BDNF-treated neurons were significantly reduced compared to naïve. Neurons sampled: ‘silent’ n = 20 (849 fluctuations), naïve n = 78 (2651 fluctuations), and BDNF n = 57 (1555 fluctuations). Median with quartiles shown. Comparisons made using Brown-Forsythe and Welch’s ANOVA tests and Games-Howell post-hoc test. **(E)** Relative frequency (%) of the fluctuations binned by amplitude. Amplitude values were normalized to the largest fluctuation in each neuron. **(F-G)** The mean fluctuation amplitude and AUC in the naïve and BDNF-treated neurons were significantly increased compared to the ‘silent’ neurons. **(H-J)** Further analysis revealed little to no effect of BDNF on fluctuation duration or frequency, but the fluctuations in ‘silent’ neurons exhibited longer rise times. For scatterplots F-J, the fluctuations that were <10% the maximal fluctuation amplitude in each neuron were omitted to reduce the effects of low-amplitude noise. Scatterplots in F-J show the median with the interquartile range. All comparisons of medians in F-K were made using Kruskal-Wallis tests and Dunn’s post-hoc test. \*\**P* < 0.01, \*\*\**P* < 0.001, \*\*\*\**P* < 0.0001.

**Table 1.**
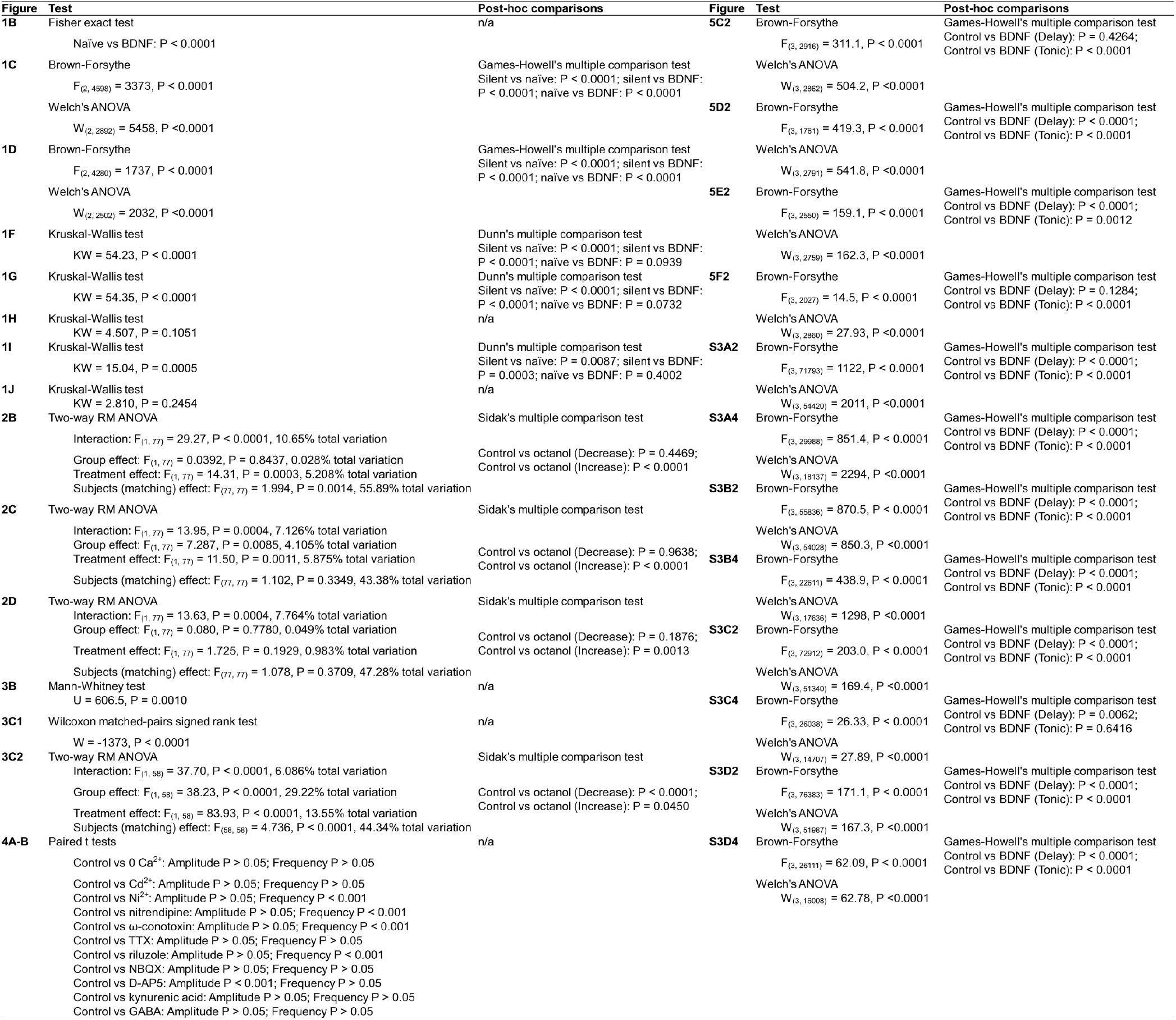
Detailed results of statistical analysis per figure.

### Octanol, a gap junction blocker, disinhibits a population of superficial dorsal horn neurons in BDNF-treated cultures

It has been shown that gap junctions have regulatory control over network activity in the *substantia gelatinosa* (23–26). Therefore, gap junctions may play a role in regulating the properties of [Ca2+]_i_ across the network, but the response of individual neurons to the chronic BDNF in the culture may be masked by the complex responses of the different types of neurons to BDNF (20,27). To address this and determine the potential role of gap junctions in mediating the BDNF-induced fluctuations, we recorded [Ca2+]_i_ from SDH neurons exposed to chronic BDNF and then applied a non-specific gap junction blocker octanol at 1mM for ≥3 min. We again performed a detailed analysis using the FIBSI program and a <10% maximal fluctuation amplitude cut-off was applied to each neuron. Inspection of the FIBSI-processed recordings revealed octanol disproportionately affected some neurons compared to others (i.e., large increases in amplitude and AUC in some neurons vs minor decreases in others). Neurons were grouped based on whether the octanol treatment caused a negative (decrease, n = 39) or positive (increase, n = 40) fold change in mean fluctuation amplitude compared to the BDNF (control) condition (**Fig 2A**). Paired analyses indicated blocking gap junctions with octanol decreased the frequency of the BDNF-induced fluctuations (**Fig 2B**) while increasing their duration (**Fig 2C**) only in the neurons that had a positive fold change in mean fluctuation amplitude. Fluctuation rise time was also significantly decreased in the same group of neurons (**Fig 2D**). These data suggest that blocking gap junctions with octanol selectively disinhibited the response to chronic BDNF in a subpopulation of SDH neurons.

**Figure 2.**
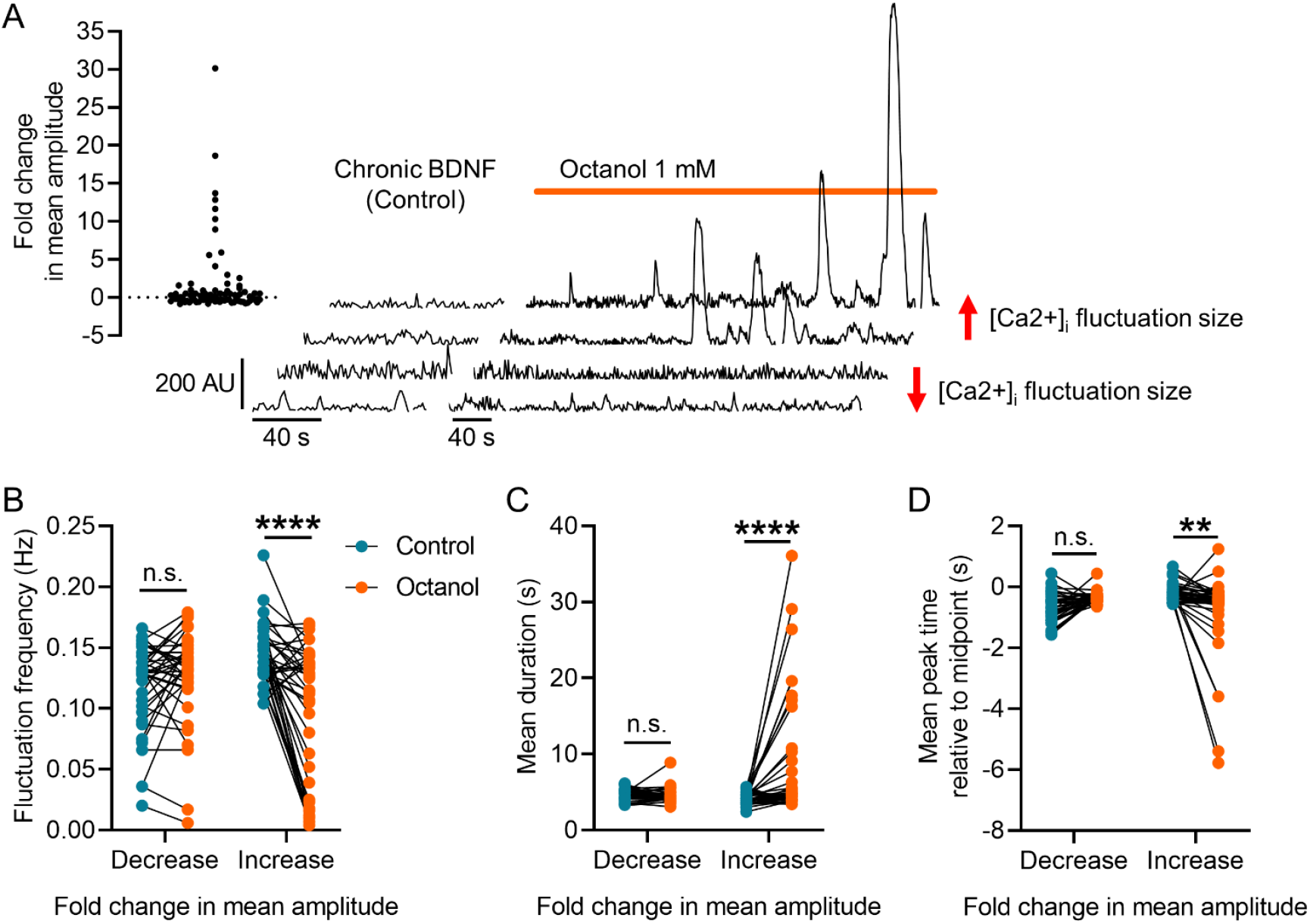
Effect of gap junction blocker octanol on BDNF-induced [Ca2+]_I_ fluctuations. **(A)** Scatterplot summarizing the effects of octanol on mean fluctuation amplitude and examples of FIBSI-processed recordings. The two bottom recordings show a decrease in fluctuation amplitude and the two top show an increase. **(B-D)** Neurons were sorted based on whether they exhibited a positive or negative fold change in mean fluctuation amplitude in response to octanol. Paired analyses showed octanol selectively and significantly decreased the mean fluctuation frequency, increased the mean fluctuation duration, and decreased the mean fluctuation rise time (i.e., more negative peak time) in the neurons with positive fold changes in mean fluctuation amplitude. Control vs octanol paired comparisons for the groups in B-D were made using two-way repeated measures ANOVAs and Sidak’s post-hoc test. \*\**P* < 0.01, \*\*\*\**P* < 0.0001.

### Chronic BDNF induces synchrony of [Ca2+]_i_ fluctuations in organotypic cultures, and blocking gap junctions reduces BDNF-induced synchrony

Previous work has shown BDNF-treated SDH neurons generate synchronous [Ca2+]_i_ fluctuations (20). We verified that finding by using the FIBSI-processed recordings and comparing the degree of synchrony between neurons imaged together in naïve and BDNF-treated cultures. Adjacency matrixes were generated using Pearson *r* coefficients; the recording of each neuron was correlated with every other neuron imaged within the same naïve or BDNF-treated culture using a Pearson correlation matrix. The adjacency matrixes for the BDNF-treated cultures showed the neurons were more positively correlated with Pearson *r* coefficients closer to 1 (**Fig 3A**), suggesting they exhibited more synchronous [Ca2+]_i_ fluctuations than the naïve cultures. To confirm this prediction, the mean coefficient for each neuron was calculated (i.e., the mean correlation of each neuron with every other neuron in its respective culture; refer to Fig 3A for Fisher Z transformation steps). Comparison of the mean Pearson *r* coefficients revealed BDNF-treated neurons were more positively correlated with other neurons in the culture than naïve neurons (**Fig 3B**). Considering the role gap junctions may play in regulating network activity and synchronous activity, this result led us to anticipate that blocking gap junctions may reduce BDNF-induced synchrony within the culture. We used the same approach to measure synchrony among neurons within the same cultures treated with BDNF prior to application of octanol, and then we asked whether blocking gap junctions caused a decrease in the mean Pearson *r* coefficients. Indeed, octanol caused a net decrease in the mean Pearson *r* coefficient (**Fig 3C1-2**). Closer inspection revealed octanol decreased the mean Pearson *r* coefficient in 100% (25/25) of the neurons in the group with decreased fluctuation amplitudes, while only 63% (22/35) of the neurons in the group with increased fluctuation amplitudes exhibited a decreased mean Pearson *r* coefficient. This suggests that blocking gap junctions may actually increase [Ca2+]_i_ fluctuation synchrony in some neurons.

**Figure 3.**
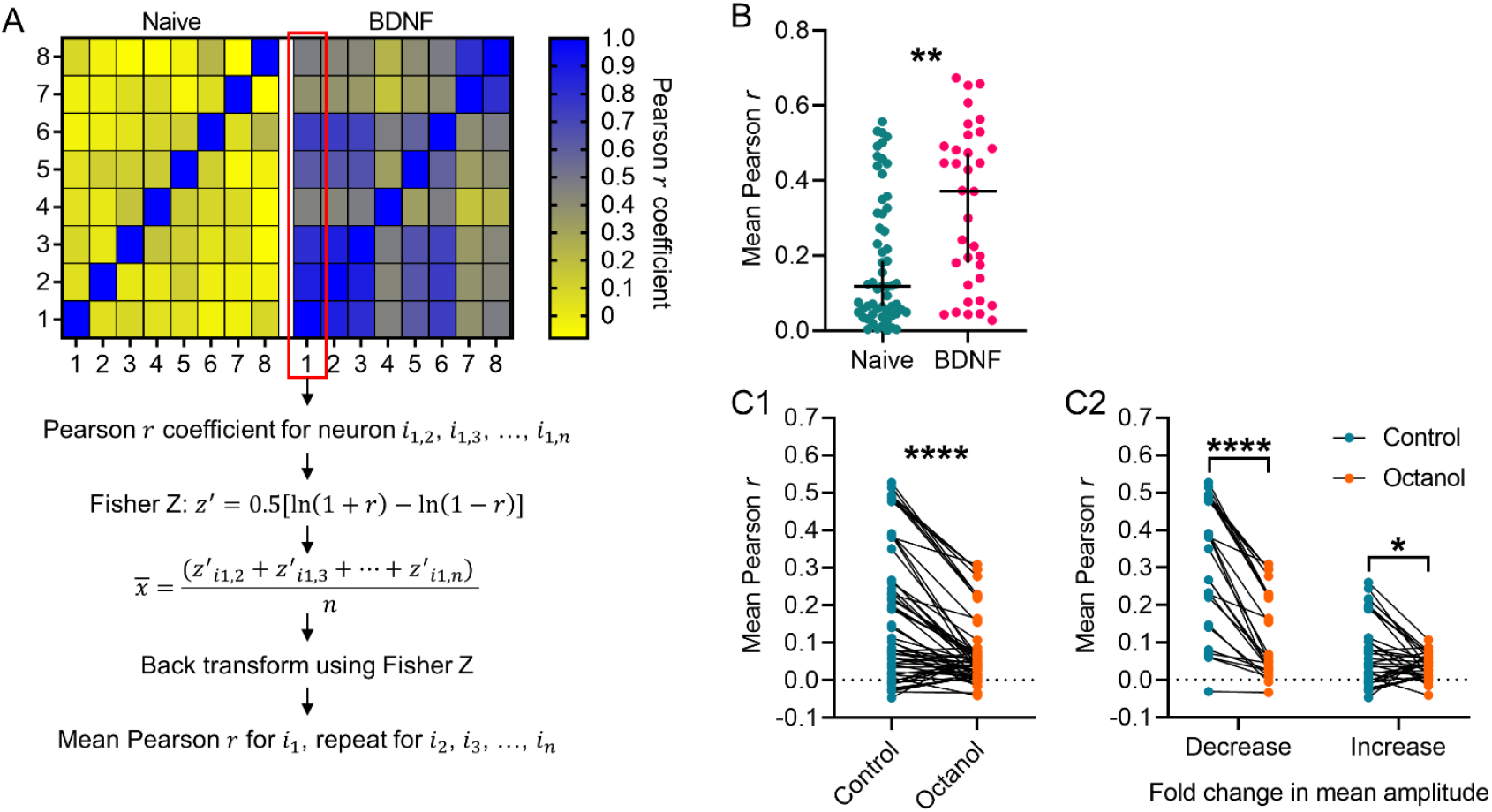
Synchrony of [Ca2+]_i_ fluctuations in BDNF-treated cultures and the effect of octanol. **(A)** Example adjacency matrix constructed from Pearson *r* coefficients corresponding to the correlation in activity between neurons in a naïve or BDNF-treated culture, each with 8 sampled neurons. Steps to calculate the mean Pearson *r* for each neuron within a culture using the Fisher Z transformation are also depicted. **(B)** Neurons sampled from BDNF-treated cultures exhibited significantly greater mean Pearson *r* coefficients compared to neurons sampled from naïve cultures. Neurons were imaged from 4 naïve cultures and 3 BDNF-treated cultures, each with ≥8 neurons imaged. Medians shown with the 95% confidence interval. Medians were compared using a Mann-Whitney test. **(C1-C2)** Treatment with octanol significantly decreased the mean Pearson *r* in neurons sampled from cultures exposed to chronic BDNF, and this treatment effect impacted both subsets of neurons binned based on their initial response to octanol. Neurons were imaged from 4 cultures. The control vs octanol paired comparison in C1 was made using a Wilcoxon rank sum test. Control vs octanol paired comparisons for the groups in C2 were made using a measures two-way repeated measures ANOVA and Sidak’s post-hoc test. \**P* < 0.05, \*\**P* < 0.01, \*\*\*\**P* < 0.0001.

### Pharmacology of [Ca2+]_i_ fluctuations in BDNF-treated slices

In order to further characterize the properties of the [Ca2+]_i_ fluctuations, we used pharmacological treatments to block a variety of voltage-gated ion channels, glutamate receptors, and GABA receptors. Comparisons between the pharmacologically-treated neurons and the BDNF-treated (control) neurons revealed the [Ca2+]_i_ fluctuation amplitudes (**Fig 4A**) and frequency (**Fig 4B**) are controlled by diverse mechanisms (example recordings and data available in **Supplemental Figure 1**). The fluctuations were ablated when extracellular Ca2+ was removed, when all voltage-gated calcium channels (VGCCs) were blocked with Cd2+, when all TTX-sensitive voltage-gated sodium channels (VGSCs) were blocked with tetrodotoxin (TTX), when AMPA or kainite glutamate receptors were blocked, and when GABA was applied to the cultures. Interestingly, the fluctuations remained when we blocked T-type (with Ni2+), L-type (with nitrendipine), or N-type (with ω-conotoxin) Ca2+ channels, although the frequency was significantly reduced with no effect on amplitude. Riluzole, which blocks glutamate release and VGSCs, only affected the fluctuation frequency but not amplitude. Finally, the NMDA blocker D-AP5 was the only blocker to significantly reduce fluctuation amplitudes, but it had no effect on frequency. These data suggest the BDNF-induced [Ca2+]_i_ fluctuations are controlled by a combination of VGCCs, TTX-sensitive VGSCs, GABA and NMDA receptors.

**Figure 4.**
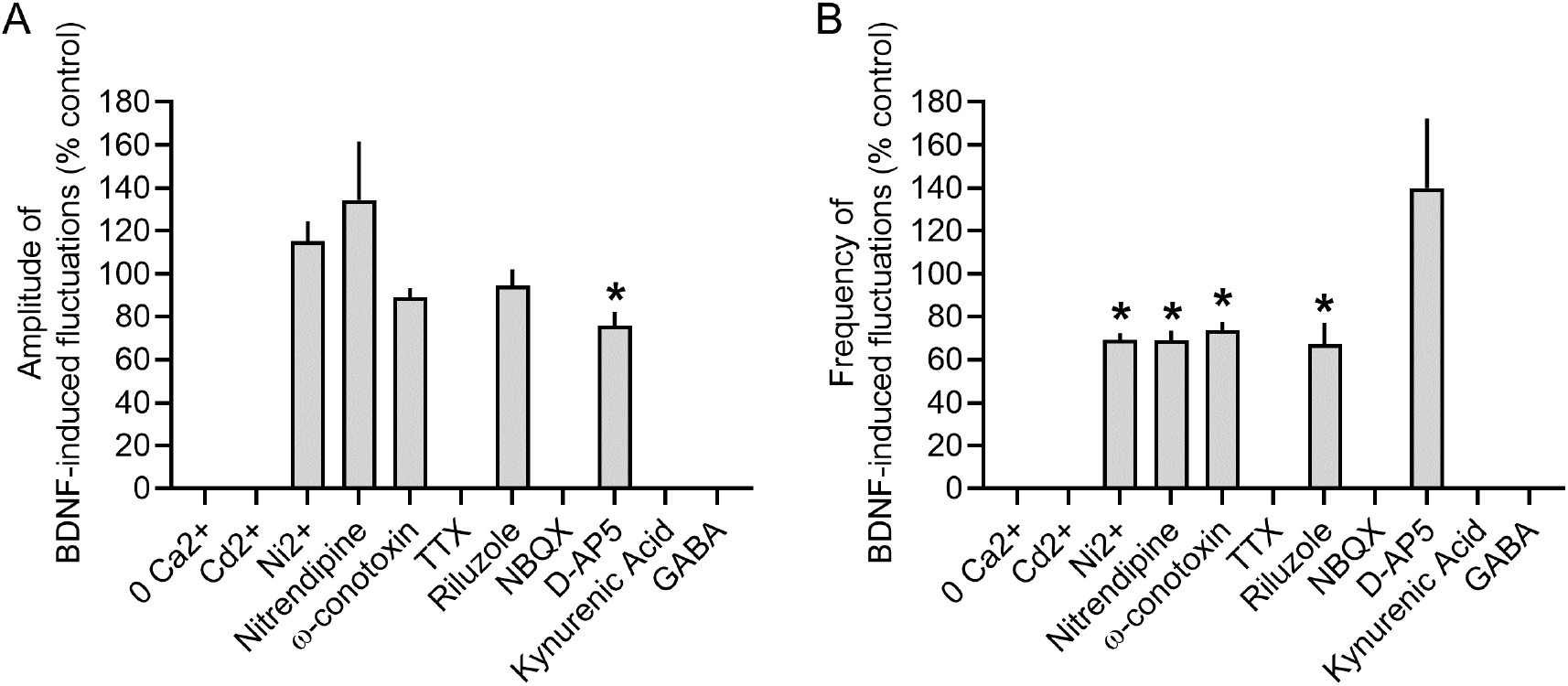
Pharmacology of [Ca2+]_i_ fluctuations from BDNF-treated cultures. Summary of the effects of various pharmacological treatments on the **(A)** amplitude and **(B)** frequency of the [Ca2+]_i_ fluctuations in BDNF-treated cultures compared to before drug treatment. Bars show the mean + standard error of the mean. \**P* < 0.001, paired t-test, n = 5-20 cells per condition.

### Clustering spontaneous excitatory postsynaptic currents (sEPSCs) reveals chronic BDNF treatment depresses excitability of delay and tonic-firing superficial dorsal horn neurons, mirroring the effect of BDNF on [Ca2+]_i_ fluctuations

It is well established that BDNF release in the SDH is involved in neuropathic pain (10,16,17). However, to our knowledge our findings are the first to show chronic BDNF treatment unmasks [Ca2+]_i_ fluctuations in a subpopulation of SDH neurons while depressing activity in others. We had previously shown that sEPSCs recorded from SDH neurons are perturbed in BDNF-treated spinal cord organotypic cultures (20). It is plausible that the newly discovered unmasked BDNF-induced [Ca2+]_i_ fluctuations underlie changes in network excitability. We decided to readdress the effects of BDNF on network excitability by using the FIBSI program to quantitatively analyze a previous subset of sEPSC recordings in delay and tonic-firing SDH neurons (example recordings and low-pass filtering/matching methods are available in **Supplemental Figure 2**). Inspection of the sEPSC interevent interval data indicated there might be natural clustering between single “small” sEPSCs and larger summated sEPSCs. We used partitioning around medoids (a medoid is like a centroid, but is restricted to an actual observation in the dataset) to cluster the sEPSCs in each neuron based on 3 sEPSC parameters: interevent interval (ms), amplitude (pA), and charge transfer (*Q*; essentially area under the curve). Optimal clustering (k medoids = 3-5 per neuron) identified 3 main clusters of sEPSCs that were present in all delay (**Fig 5A1-A2**) and tonic (**Fig 5B1-B2**) neurons in both control and BDNF treatment conditions (clusters named based on amplitude; interevent interval): 1) small; short, 2) small; long, and 3) large. Two other clusters were identified (small; mid and medium), but they were not present in all neurons or treatment conditions and were not analyzed for this study. We were mainly interested in the effects of BDNF on the cluster of large-amplitude sEPSCs in both neuron types (refer to **Supplemental Figure 3** for the “small” sEPSC analysis). Chronic BDNF treatment significantly decreased sEPSC amplitudes (**Fig 5C1-C2**) and charge transfer (**Fig 5D1-D2**) in the tonic-firing neurons. We did not observe a significant effect on sEPSC amplitudes in the delay neurons, but charge transfer was significantly decreased. The sEPSCs in the BDNF-treated delay neurons were narrower (control duration = 83.6 ± 1.2 ms; BDNF duration = 49.9 ± 0.4 ms). The sEPSC interevent intervals were significantly shorter in both neuron types (**Fig 5E1-E2**), and peak time measurements indicated BDNF significantly increased activation of the sEPSCs in the tonic neuron (**Fig 5F1-F2**).

**Figure 5.**
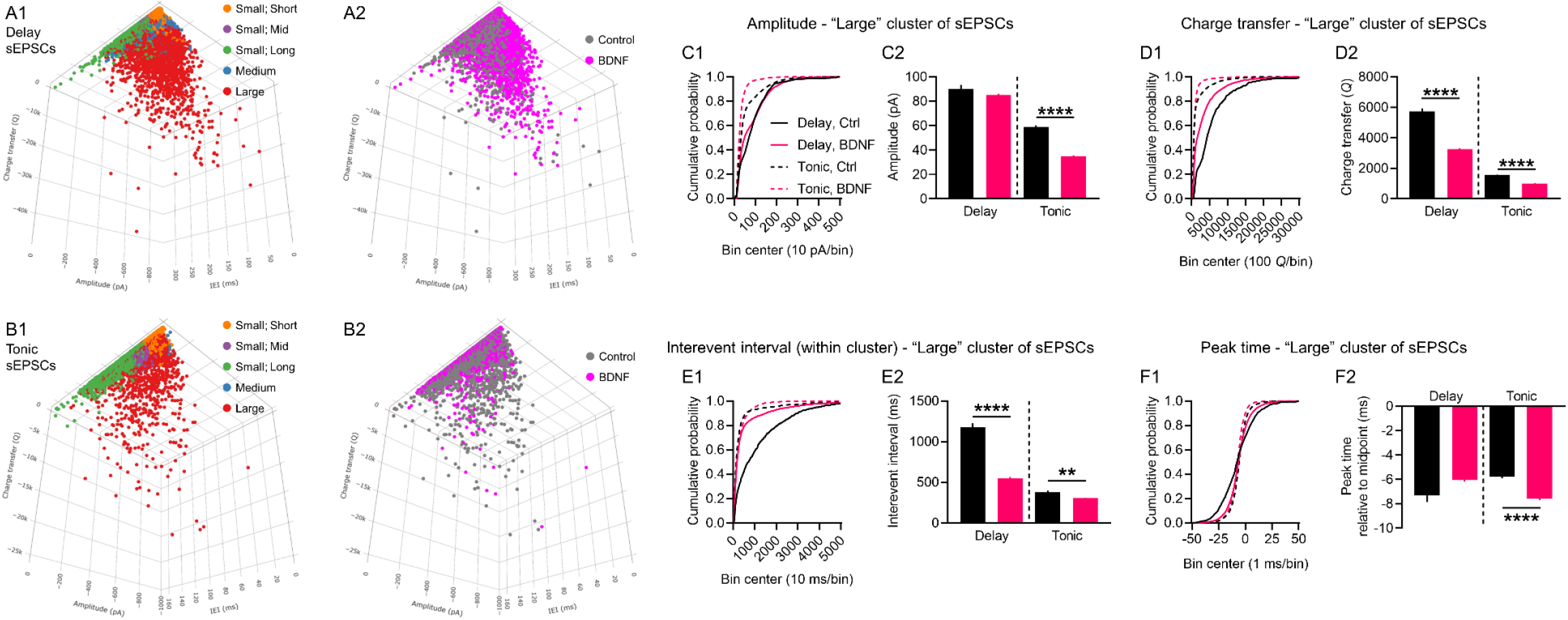
Cluster analysis of spontaneous EPSCs in delay and tonic-firing superficial dorsal horn neurons. Three dimensional representations of the sEPSCs in **(A1-A2)** delay neurons and **(B1-B2)** tonic neurons plotted based on interevent interval (x-axis), amplitude (y-axis), and charge transfer (z-axis). Mapping of the clustering results and treatment shown for the total sample of sEPSCs. The sEPSCs in each neuron were clustered independently from the other neurons. Optimal partitioning around medoids identified 3 primary clusters present in all neurons in both treatment conditions (naming based on amplitude; interevent interval): small; short, small; long, and large. Two additional clusters were identified in some neurons, but were not present in all neurons in both conditions: small; mid and medium. Neurons sampled: 19 delay (5 control, 14 BDNF); 9 tonic (5 control, 4 BDNF). Total number of sEPSCs: 33,206 in delay, control; 108261 in delay, BDNF; 40658 in tonic, control; 36521 in tonic, BDNF. The effects of BDNF on sEPSC amplitude, charge transfer (i.e., area under the curve), and peak time were assessed for the 3 primary clusters (large cluster shown, refer to Supplemental Figure 3 for the two small-amplitude clusters). **(C1-C2)** Treatment with BDNF significantly decreased sEPSC amplitudes in tonic neurons, but a significant effect was not observed in the delay neurons. Cumulative distributions of the measured amplitudes shown (left) alongside the means ± standard error of the means (right). **(D1-D2)** The BDNF treatment significantly reduced sEPSC charge transfer measures in both delay and tonic neurons. **(E1-E2)** Interevent intervals for the sEPSCs within the large cluster were significantly decreased in the BDNF-treated neurons. **(F1-F2)** The sEPSC activation kinetics were significantly faster in the BDNF-treated neurons, but no significant effect was observed for the sEPSCs in the delay neurons. Large sEPSC cluster sample sizes: delay neurons control n = 753, BDNF n = 4539; tonic neurons control n = 2348, BDNF n = 2367. Comparisons between means were made using Brown-Forsythe and Welch’s ANOVA tests and Games-Howell post-hoc test. \*\**P* < 0.01, \*\*\*\**P* < 0.0001.

## Discussion

The present study was undertaken to better understand mechanisms underlying synchronous activity among SDH neurons and the influence BDNF may have on network excitability in naïve, uninjured organotypic cultures. Our experiments indicate long-term exposure to BDNF produces a complex set of neuron-dependent changes in network excitability in the SDH. First, BDNF causes a concomitant activation of Ca2+ signaling in a subpopulation of ‘silent’ SDH neurons (∼20% sampled) and depression of Ca2+ signaling in many others. This finding was largely unexpected as we have previously shown chronic BDNF increases the size and frequency of [Ca2+]_i_ fluctuations (20). It is plausible that the presence of the ‘silent’ neurons can greatly influence data analysis and interpretation depending on how they are grouped together, or discarded. Another possibility is that resolution differences between the two event-detection programs used (FIBSI in the current study, Mini Analysis Software (Synaptosoft, NJ)) in the previous study) could influence analysis of the [Ca2+]_i_ fluctuations. However, the goal of the present study was not to compare/contrast the two programs. The second significant finding was that gap junction signaling may play a crucial role in regulating BDNF-induced changes in Ca2+ signaling in the SDH. Blockade of gap junctions with octanol unveiled a subpopulation of neurons that respond to BDNF with large-amplitude, low-frequency [Ca2+]_i_ fluctuations. This finding suggests that gap junctions modulate global network activity, either by directly or indirectly inhibiting those particular neurons with increased Ca2+ signaling. Indeed, this is supported by the third major finding that BDNF profoundly increases synchrony of the [Ca2+]_i_ fluctuations, and blocking gap junctions with octanol reverses BDNF-induced synchrony.

While the physiological correlates of the BDNF-related [Ca2+]_i_ fluctuations are unclear, we do not think they are being driven by delay or tonic-firing neurons in the SDH. Unlike our analysis of the [Ca2+]_i_ fluctuations where we describe the ‘silent’ neurons, we did not observe any unmasking effect(s) of BDNF on the sEPSCs in the delay or tonic neurons. All neurons sampled in the control (naïve) condition generated visually appreciable, large-amplitude sEPSCs. The main depressive effect of BDNF on the sEPSCs (refer also to **Supplemental Figure 3**) mirrored the main effect of BDNF on comparable properties for the [Ca2+]_i_ fluctuations. A summary of each dataset (**Fig 6**, table) shows BDNF reduced the amplitude and area under the curve parameters for the [Ca2+]_i_ fluctuations and large sEPSCs. More nuanced effects were observed for frequency (reciprocal of interevent interval) and activation kinetics: BDNF increased the frequency of the large sEPSCs but had little to no effect on the frequency of the [Ca2+]_i_ fluctuations; BDNF increased activation of the large sEPSCs in the tonic neurons and had little to no effect on the delay neurons or [Ca2+]_i_ fluctuations. However, these data collectively suggest chronic BDNF may have reduced excitability in the delay and tonic SDH neurons, and this may provide a physiological basis for the depressive effect of chronic BDNF on the [Ca2+]_i_ fluctuations. On the contrary, the ‘silent’ neurons are predicted to have responded to chronic BDNF with [Ca2+]_i_ fluctuations of greater amplitude, total area, frequency, and presumably activation. Our working model (**Fig 6**, bottom) posits that under naïve conditions many delay and tonic SDH neurons in organotypic spinal cord cultures are generally in an active state, with some gap junction coupling with neighboring neurons. Long-term exposure to BDNF causes a homeostatic shift that pushes the SDH into a dampened state marked by synchronous, low-level activity spread across the network via increased gap junction coupling. This synchronous activity in turn mitigates the BDNF-activated ‘silent’ neurons.

**Figure 6.**
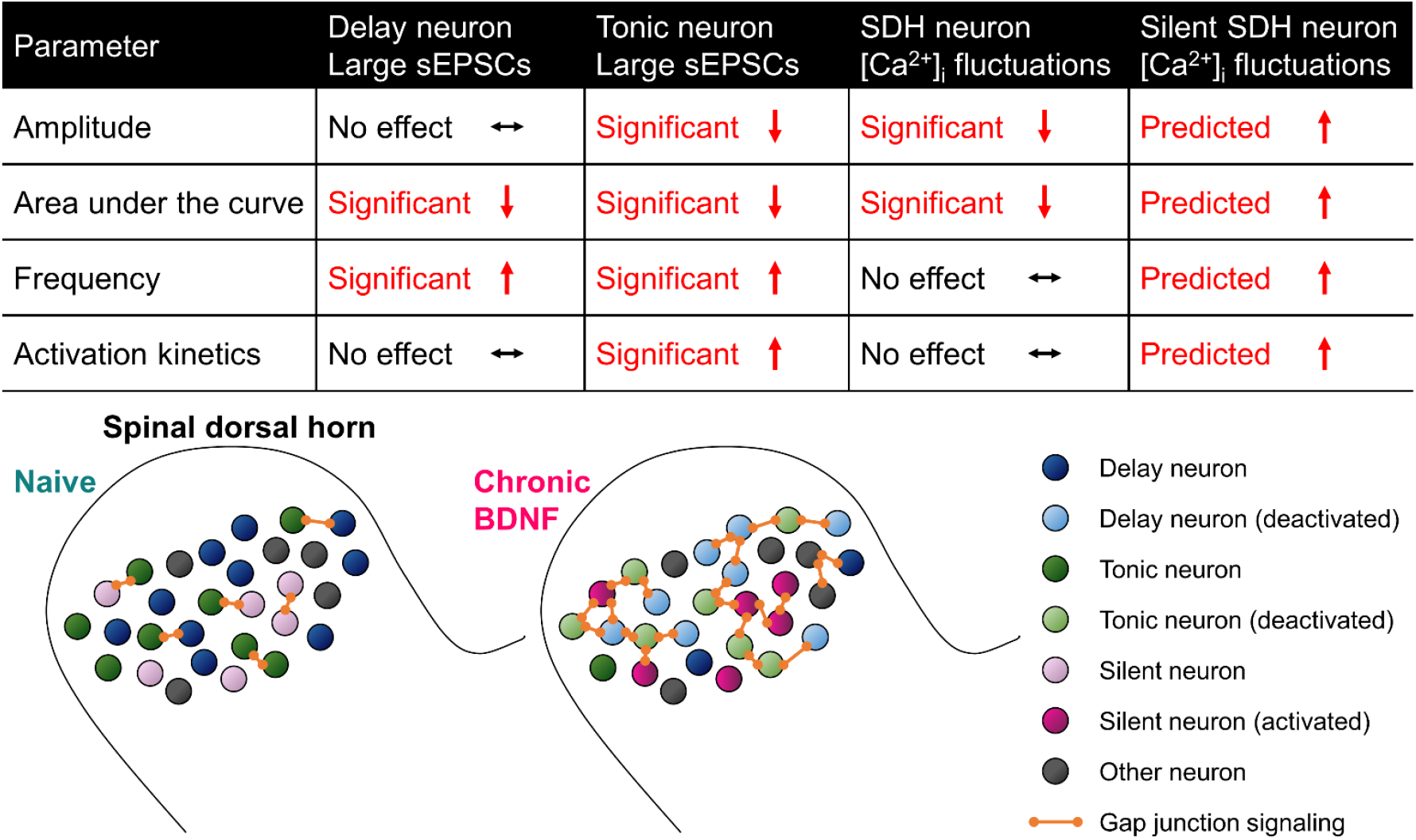
Working model summarizing the effects of chronic BDNF on SDH neurons in this study.

Several types of oscillatory and/or rhythmic bursting activity have been previously observed in spinal cord organotypic cultures (28,29) and in *ex vivo* slice preparations (23,30–32). Synchronous activation of groups of spinal nociceptive neurons might contribute to the ‘electric shock’-like sensations experienced by some neuropathic pain patients (33). We hypothesize that [Ca2+]_i_ fluctuations leading to synchronous network activity may be attributed to the “shooting pains” that chronic pain patients may experience. In particular, “shooting pain” is very commonly experienced by patients with radicular lower back pain (sciatica) or trigeminal neuralgia. In these patients, stimulus movement elicits so-called “traveling waves” in which neural activity sweeps across the body-part representation in somatosensory maps (34). The mechanisms of this have been attributed in part to Ca^2+^ waves and gap junctions (34,35). Therefore, given the properties of the [Ca2+]_i_ fluctuations we have identified, they would make a good candidate for the neural substrate of shooting pain. In a cohort of chemotherapy-induced peripheral neuropathy (CIPN) patients, some of whom reported shooting or burning pain in the hands or feet, serum nerve growth factor (NGF) levels were much higher compared with patients with painless or absent CIPN (36). Therefore, there is a precedent for the correlation of neurotrophic factors related to BDNF to the incidence of shooting pain. Indeed it has been shown that NGF regulates the expression of BDNF (37). It is has also been demonstrated that BDNF overexpression induces spasticity in rodent models of spinal cord injury, which may also explain the effect of BDNF on influencing network excitability in the dorsal horn (38). With regards to the involvement of the dorsal horn in shooting pain, it has been shown that dorsal root entry zone lesioning (DREZotomy) successfully reduces shooting pain caused by brachial plexus root avulsion (BPRA) (39). Also, in one patient with shooting pain caused by osteoid osteoma, a partial laminectomy was able to significantly relieve pain symptoms (40). Ideally however, more experiments using human *ex vivo* tissue are needed to determine whether these particular [Ca2+]_i_ fluctuations correlate with shooting pain in patients.

## Materials and Methods

### Defined-medium organotypic cultures of spinal cord slices

Spinal cords were isolated from embryonic (E13-14) rats and transverse slices (300 ± 25 μm) were cultured using the roller-tube technique (20,41). Since tissue is obtained from embryos, sex could not be determined. Serum-free conditions were established after 5 days *in vitro*. Medium was exchanged with freshly prepared medium every 3-4 days. Slices were treated after 15-21 days *in vitro* for a period of 5-6 days with 50-200 ng/ml in serum-free medium as described previously (20). Age-matched, untreated DMOTC slices served as controls.

### Calcium imaging

Each organotypic slice was incubated for 1 h prior to imaging with the fluorescent Ca^2+^-indicator dye Fluo-4-AM (Molecular Probes, Invitrogen, Carlsbad, CA, USA). The conditions for incubating the dye were standardized across different slices to avoid uneven dye loading. After dye loading, the slice was transferred to a recording chamber and superfused with external solution containing (in mM): 131 NaCl, 2.5 KCl, 1.2 NaH_2_PO_4_, 1.3 MgSO_4_, 26 NaHCO_3_, 25 D-glucose, and 2.5 CaCl_2_ (20°C, flow rate 4 ml/min). Regions of interest (ROI) corresponding to individual cell bodies of neurons were identified based on morphology and size. Changes in Ca^2+^-fluorescence intensity evoked by a high K^+^ solution (20, 35, or 50 mM, 90 s application)or other pharmacological agents, were measured in dorsal horn neurons with a confocal microscope equipped with an argon (488 nm) laser and filters (20x XLUMPlanF1-NA-0.95 objective; Olympus FV300, Markham, Ontario, Canada). Full frame images (512 × 512 pixels) in a fixed xy plane were acquired at a scanning time of 0.8-1.08 s/frame (42). In some experiments, images were cropped to accommodate faster scan rates. Selected regions of interest were drawn around distinct cell bodies and fluorescence intensity traces were generated with FluoView v.4.3 (Olympus).

### Electrophysiology

Whole cell patch-clamp recordings were obtained from neurons in organotypic slice cultures under infrared differential interference contrast optics. Neurons selected for recording were located 250–800 μm from the dorsal edge of the cultures in an area presumed to reflect the *substantia gelatinosa* and up to a depth of 100 μm from the surface. Neurons were categorized according to their firing pattern in response to depolarizing current commands as tonic, delay, phasic, irregular, or transient (27). Recordings were obtained with an NPI SEC-05LX amplifier (ALA Scientific Instruments, Westbury, NY, USA) in bridge balance or in discontinuous, single electrode, current or voltage-clamp mode. Neurons were sampled at 2k Hz for 180 seconds when measuring sEPSCs. For recording, slices were superfused at room temperature (∼22°C) with 95% O2-5% CO2-saturated aCSF that contained (in mM) 127 NaCl, 2.5 KCl, 1.2 NaH2PO4, 26 NaHCO3, 1.3 MgSO4, 2.5 CaCl2, and 25 d-glucose, pH 7.4. Patch pipettes were pulled from thin-walled borosilicate glass (1.5/1.12 mm OD/ID; WPI, Sarasota, FL) to 5-to 10-MΩ resistances when filled with an internal solution containing (in mM) 130 potassium gluconate, 1 MgCl2, 2 CaCl2, 10 HEPES, 10 EGTA, 4 Mg-ATP, and 0.3 Na-GTP, pH 7.2, 290–300 mosM.

### Drugs and chemicals

Unless otherwise stated, all chemicals were from Sigma (St. Louis, MO, USA). Fluo-4 AM dye was dissolved in a mixture of dimethyl sulfoxide (DMSO) and 20% pluronic acid (Invitrogen, Burlington, Ontario, Canada) to a 0.5 mM stock solution and kept frozen until used. The dye was thawed and sonicated thoroughly before incubating with a DMOTC slice. TTX was dissolved in distilled water as a 1 mM stock solution and stored at -20°C until use. TTX was diluted to a final desired concentration of 1 µM in external recording solution on the day of the experiment. Strychnine was prepared in a similar manner to TTX, and bicuculline (Tocris, Ballwin, MO, USA) was dissolved in DMSO as a 10 mM stock solution. 6-cyano-7-nitroquinoxaline-2,3-dione (CNQX, Tocris), 2,3-dihydroxy-6-nitro-7-sulfamoyl-benzo[f]quinoxaline-2,3-dione (NBQX, Tocris) and N,N,H,-Trimethyl-5-[(tricyclo[3.3.1.13,7]dec-1-ylmethyl)amino]-1-pentanaminiumbromide hydrobromide (IEM-1460, Tocris) were prepared as 10 mM stocks dissolved in distilled water, and D(-)-2-Amino-5-phosphonopentanoic acid (D-AP5, Tocris) was prepared as 50 mM stocks dissolved in 30% 1 M NaOH. Riluzole was prepared as a 10 mM stock, kyneurinic acid and GABA as 1 M stocks, and nitrendipine as a 1 mM stock made up in distilled water. These drugs were used at a 1:1000 dilution prepared freshly with external recording solution immediately prior to the start of experiments.

### Data analysis and statistical testing

Raw time-lapse calcium fluorescence intensity and whole-cell voltage-clamp recordings were analyzed using the Frequency-Independent Biological Signal Identification (FIBSI) program (21) written using the Anaconda v2019.7.0.0 (Anaconda, Inc, Austin, TX) distribution of Python v3.5.2 and the NumPy and matplotlib.pyplot libraries. The FIBSI program incorporates the Ramer-Douglas-Peucker algorithm to detect significant waveforms against a time-dependent generated reference line using the least number of total points to represent said waveform, and provides quantitative measurements for each detected waveform. The calcium fluorescence traces were fit using a sliding median with a window size between 5-25 s, and peaks below the sliding median line were traced together to form the reference line. The voltage-clamp traces were fit using a sliding median with a window of 50 ms, and peaks above the sliding median were traced together to form the reference line. A custom Python script was used to match sEPSCs detected in filtered voltage-clamp recordings with the unfiltered recordings. The detected sEPSCs for each neuron were clustered based on their amplitude, charge transfer, and interevent interval using the partitioning around medoids function (default settings, manhattan distance used) in the cluster v2.1.0 package in R v4.0.2 (2020-06-22; R Core Team, The R Foundation for Statistical Computing, Vienna, Austria). Each neuron was clustered independently. The average silhouette width method was used to select the optimal number of clusters for each neuron. Clusters were graphed using the plotly v4.9.2.1 package with dependency on ggplot2. The FIBSI source code, custom Python matching script, and a tutorial for using FIBSI are available on a GitHub repository (https://github.com/rmcassidy/FIBSI_program).

Statistical analysis of the calcium fluctuations and sEPSCs were performed using Prism v8.2.1 (GraphPad Software, Inc, La Jolla, CA). The comparison between the proportion of ‘silent’ neurons in the naïve and BDNF treatment conditions was made using Fisher’s exact test. The Brown-Forsythe and Welch ANOVA tests and Games-Howell post-hoc test (for n > 50) were used to compare the effects of BDNF when the calcium fluctuations from all neurons were grouped together. To control for differences in recording times, the means for each fluctuation parameter (amplitude, AUC, etc.) were calculated for each neuron (fluctuations <10% the maximal amplitude in each neuron were omitted to remove the effects of low-amplitude noise). The amplitude and AUC means were normalized across all neurons to permit direct comparisons. Normality was assessed using the D’Agostino & Pearson omnibus test, and nonparametric comparisons between group medians were made using the Kruskal-Wallis test and Dunn’s post-hoc test. Two-way repeated measures ANOVAs and Sidak’s post-hoc tests were used to compare the pre- and post-treatment effects of octanol on the calcium fluctuations. Adjacency matrixes for neurons imaged together from the same organotypic culture were constructed using Pearson correlation matrixes; the FIBSI-processed calcium fluctuation recordings were used as input. Next, the mean Pearson *r* coefficient for each neuron within a culture was calculated using the Fisher Z transformation (steps shown in Fig 3A). The mean Pearson *r* coefficients in the naïve and BDNF conditions were not normally distributed, so the comparison between the two was made using a Mann-Whitney test. Comparisons between the pre- and post-treatment effects of octanol were first made using a Wilcoxon matched-pairs signed rank test, and then using a two-way repeated measures ANOVA and Sidak’s post-hoc test. Paired t-tests were used for pharmacology experiments. Brown-Forsythe and Welch ANOVA tests and Games-Howell post-hoc test were used to compare the effects of BDNF on the sEPSC clusters. Statistical significant was set to *P* < 0.05 and all reported *P* values are two-tailed. Details for the statistical analyses can be found in Table 1.

## Acknowledgements

We thank Edgar Walters for useful discussions. We thank Klaus Ballanyi and Araya Ruangkittisakul for use of and technical assistance with confocal imaging equipment.

## Supplemental Figures

**Supplementary Figure 1.**
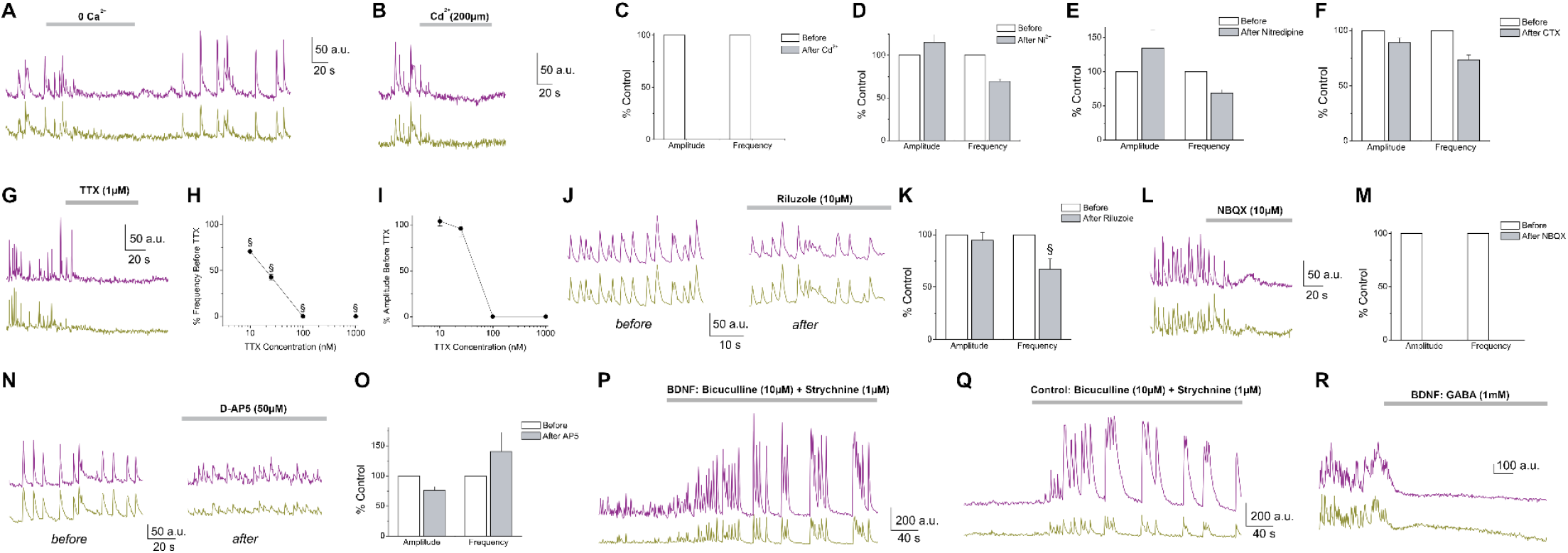
Pharmacology of BDNF-induced Ca2+ fluctuations. Dependence of BDNF-induced Ca2+ fluctuations on extracellular Ca2+ entry through voltage-gated Ca2+ channels. **(A)** Perfusion of extracellular recording solution free of Ca2+ (0 Ca2+) marked by a thick grey line. Note the complete block of the fluctuations in recorded cells. **(B)** Addition of 200 μM Cd2+, marked by a thick grey line, abolished Ca2+ oscillations in a BDNF-treated slice. **(C)** Measurement of fluctuation amplitude and frequency before and after application of Cd2+. Values normalized to control values obtained before addition of Cd2+. **(D-F)** Effect of 100 μM Ni2+, 1 μM nitrendipine, and 100 nM ω-conotoxin GVIA on BDNF-induced fluctuation amplitude and frequency. Dependence of BDNF-induced Ca2+ fluctuations on TTX-sensitive voltage-gated Na+ current but not persistent Na+ current. **(G)** Addition of 1 μM TTX, marked by a thick grey line, abolished Ca2+ fluctuations in a BDNF-treated slice. **(H)** Concentration-inhibition curve for increasing concentrations of TTX on Ca2+ fluctuation frequency. **(I)** Concentration-inhibition curve for increasing concentrations of TTX on Ca2+ fluctuation amplitude. Error bars indicate standard error of the mean. For paired t-test, § = p<0.001. **(J)** Sample synchronous Ca2+ fluctuation traces from a BDNF-treated slice before (left) and after (right) application of 10 μM riluzole. **(K)** Effect of riluzole on average Ca2+ fluctuation amplitude and frequency. BDNF-induced Ca2+ fluctuations mediated by AMPA/kainate glutamate receptors. **(L)** Addition of 10 μM NBQX, marked by a thick grey line, abolished Ca2+ fluctuations in a BDNF-treated slice. **(M)** Measurement of fluctuation amplitude and frequency before and after application of NBQX. **(N)** Sample fluorescent Ca2+ traces from a BDNF-treated slice before (left) and after (right) application of 50 μM AP5. **(O)** Effect of AP5 on average Ca2+ fluctuation amplitude and frequency. Amplification of BDNF-induced Ca2+ oscillations by pharmacological removal of inhibition and suppression of oscillatory activity by GABA. **(P)** Addition of 10 μM bicuculline and 1 μM strychnine to a BDNF-treated slice produced robust fluctuations larger in amplitude but slower in frequency than the spontaneous Ca2+ fluctuations observed before antagonist application. **(Q)** Addition of 10 μM bicuculline and 1 μM strychnine to a control DMOTC slice produced similar robust fluctuations as those observed in P. **(R)** Application of 1 mM GABA, marked by a thick grey line, stopped BDNF-induced Ca2+ fluctuations. Average values represented. Error bars indicate standard error of the mean. For paired t-test, § = p<0.001, n=5-20 cells per condition.

**Supplementary Figure 2.**
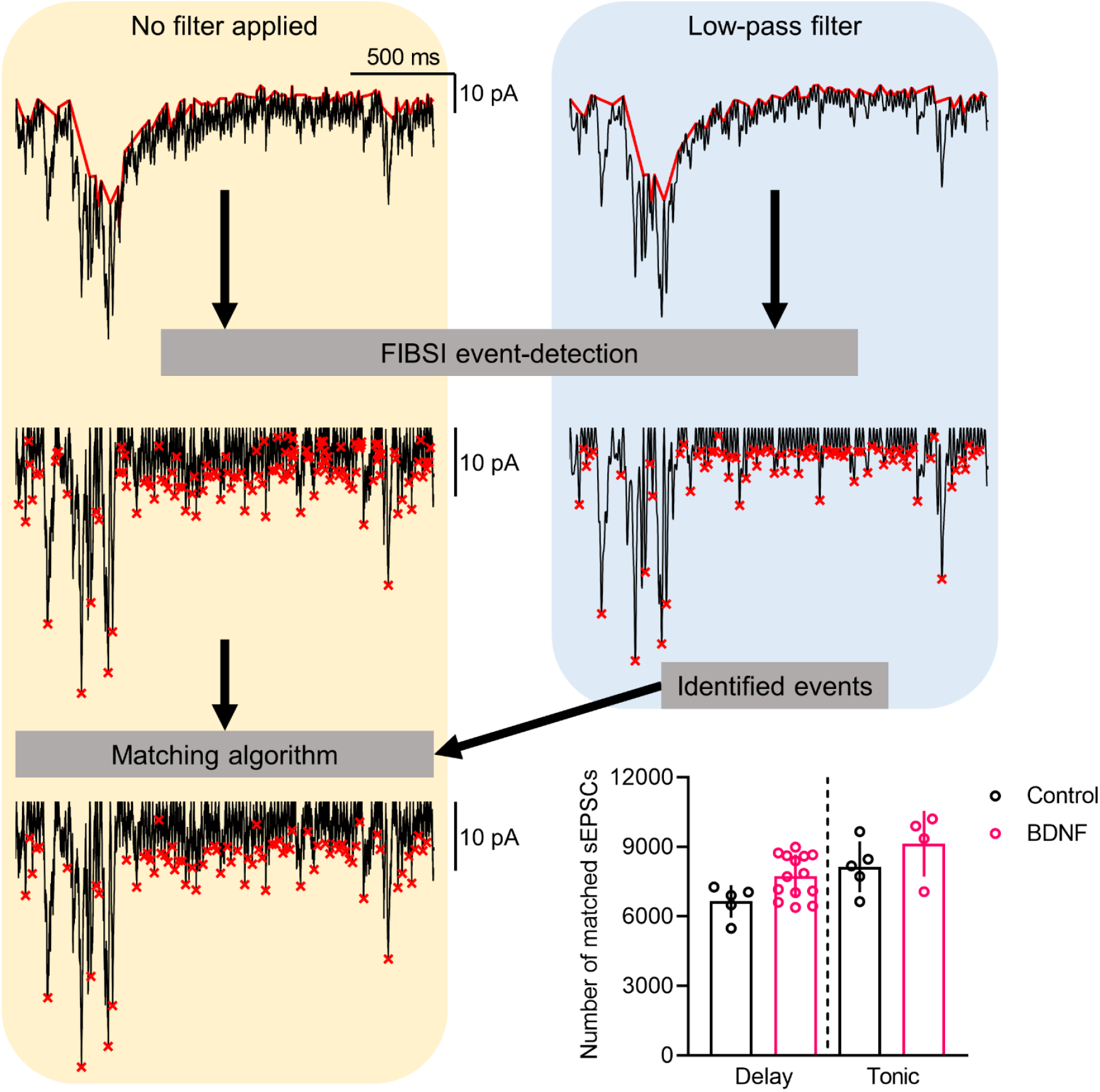
Matching filtered spontaneous EPSCs detected by FIBSI with the raw recordings. All raw recordings were first analyzed by FIBSI without pre-processing filtering. All recordings were analyzed again following application of a low-pass filter with α = 0.05. The output text files generated by FIBSI containing descriptive parameters (e.g., start time, peak time, amplitude, etc.) for each detected event were used as input to a custom Python script. The matching algorithm matched events detected in the filtered recordings with their corresponding events in the unfiltered recordings. Events were matched based on peak event time and amplitude. Unmatched events were discarded from further analyses. The total number of matched sEPSCs in the delay and tonic neurons are shown.

**Supplementary Figure 3.**
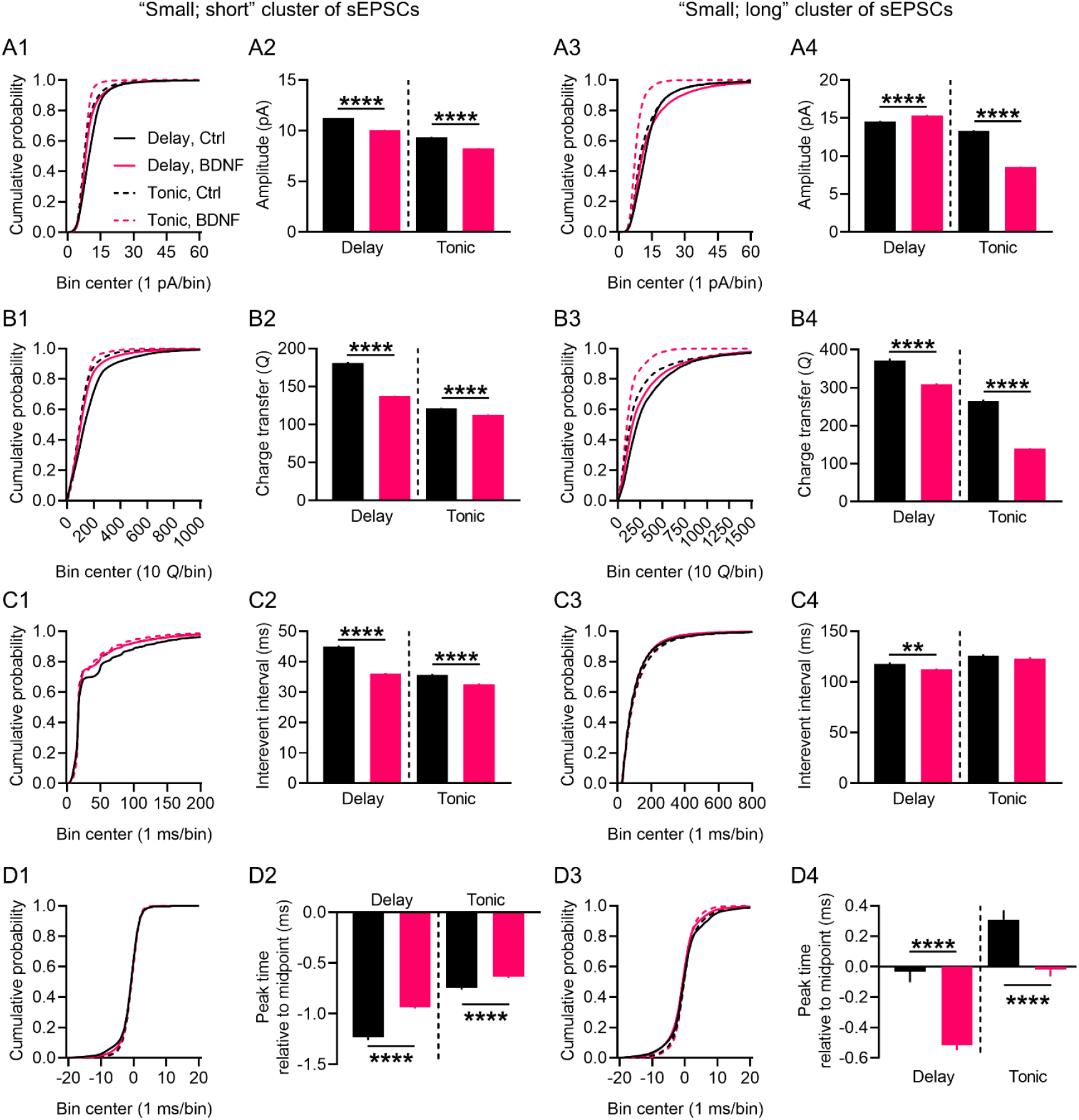
Effects of BDNF on the two small clusters of spontaneous EPSCs in delay and tonic-firing superficial dorsal horn neurons. Consistent, statistically significant effects of BDNF were observed for the ‘small; short” cluster of sEPSCs in the delay and tonic neurons; amplitudes were smaller **(A1-A2)**, charge transfer values were smaller **(B1-B2)**, within cluster interevent intervals were shorter **(C1-C2)**, and kinetics were slower **(D1-D2)**. Nuanced effects of BDNF were observed for the “small; long” cluster of sEPSCs in the two neuron types; sEPSC amplitudes were increased in delay neurons and decreased in tonic neurons **(A3-A4)** while charge transfer values were decreased in both types of neurons **(B3-B4)**. Interevent intervals in the delay neurons were significantly decreased, but not in the tonic neurons **(C3-C4)**. The sEPSC kinetics were significantly faster in the BDNF-treated delay and tonic neurons **(D3-D4)**. Small; short sEPSC cluster sample sizes: delay neurons control n = 20062, BDNF n = 69670; tonic neurons control n = 25235, BDNF n = 22086. Small; long sEPSC cluster sample sizes: delay neurons control n = 7638, BDNF n = 22391; tonic neurons control n = 7160, BDNF n = 5851. Comparisons between means in the control and BDNF conditions were made using Brown-Forsythe and Welch’s ANOVA tests and Games-Howell post-hoc test. \*\**P* < 0.01, \*\*\*\**P* < 0.0001.

